# Reliability and Validity of the Jumpster accelerometer-based app compared to the Vertec when completing a countermovement jump

**DOI:** 10.1101/2024.10.01.616202

**Authors:** Matthew E. Holman, Christopher R. Harnish

**Author notes:** All authors contributed equally to this work.

## Abstract

The reliability and validity of the Jumpster app (JA) was compared to the Vertec. Thirty-six participants completed 100 total trials using both tools simultaneously. Validity was assessed using correlation and tolerance analyses. Reliability was assessed using 95% predictive intervals (PI95) and tolerance limits (TL95) between the measures, comparing standard error of the measure (SEM) and coefficients of variation (CV) for each tool, and examining the intraclass correlation coefficient (ICC 2,K; upper and lower 95% CI) comparing both tools. The JA was weakly related to the Vertec (r = 0.24; p < 0.01). The tolerance analysis showed a moderately strong proportional bias of the JA (r = 0.45; p < 0.01). While all data fell within calculated PI95 ±TL95, the JA SEM (14.7cm) and CV (40.30%) exceeded the Vertec SEM (3.57cm) and CV (7.22%) with an ICC of 0.55 [0.79, -0.08]. These JA is neither reliable or valid.

## Introduction

Muscular power is an important aspect of health, fitness, and functional ability across the lifespan. For aging adults, the inability to produce power has been linked to an increased risk of falls and reduced functional capacity independent of other factors (Alcazar et al., 2020; Runge et al., 2004; Shur et al., 2021; Trombetti et al., 2016). Deficits in muscular power are also becoming more common among children and adolescents, resulting in impaired execution of fundamental movement skills (e.g., jumping and running) as well as a reduction in their overall health and well-being (Faigenbaum et al., 2023; Faigenbaum & MacDonald, 2017; Katartzi et al., 2005) Therefore, simple methods to assess MP across the lifespan are necessary.

The vertical jump (VJ) has long been used to measure lower extremity power, as well as neuromuscular fatigue within a variety of populations (Argaud et al., 2017; Baptista et al., 2022; Forte & Macaluso, 2008; Pupo et al., 2020; Santos et al., 2022). While most often used to assess athletic populations (Pupo et al., 2020), the VJ has also been shown to be a simple, safe, and effective measurement among children and adolescents (Baptista et al., 2022; Katartzi et al., 2005; Stojanović et al., 2020; Temfemo et al., 2009) as well as for older adults (Argaud et al., 2017; Forte & Macaluso, 2008; Santos et al., 2022). The methodology for VJ testing varies minimally, but the equipment and space needed for accurate measurement can limit its use, particularly in clinical settings. The “gold standard” tools used to assess VJ height generally require either in-ground force plates (Buckthorpe et al., 2012) and/or high-speed video analysis (Dias et al., 2011); neither of which are easily accessible in all settings or cost-conservative. The Vertec Vertical Jump Trainer (Sports Imports, Hilliard, OH, USA) has been shown to be valid and reliable when compared to both “gold standard” devices (Buckthorpe et al., 2012; Leard et al., 2007); however, it too is expensive and requires significant ceiling height for proper use. As such, low-cost tools requiring minimal equipment would broaden the use of VJ height as a clinically relevant measure of overall fitness and lower extremity power.

The advent of smart phones has resulted in a range of new fitness apps in the last decade, including apps designed to measure VJ. Arguably one of the most popular and well-studied mobile apps used to assess VJ is My Jump *2*, which is part of a suite of apps included in the My Jump Lab app. In a range of studies, various versions of this app have consistently been shown to possess high validity and reliability when compared to both the Vertec as well as electronic timing and video systems (Bogataj, Pajek, Andrašić, et al., 2020; Bogataj, Pajek, Hadžić, et al., 2020; Cruvinel-Cabral et al., 2018; Gallardo-Fuentes et al., 2016; Gençoğlu et al., 2023; Oh et al., 2020; Oliveira-Silva et al., 2018; Stanton et al., 2015; Yingling et al., 2018). However, the My Jump Lab app’s cost may be a barrier to some. Cheaper options include the *What’s my vertical* app (Dow et al., n.d.; Montalvo et al., 2021) and the JumPo 2 apps (Vieira et al., 2021, 2023), both of which have been shown to produce valid and reliable results. Compared to the My Jump 2 app, these options cost adopters considerably less money; however, like the My Jump 2 app these more economical tools also rely on the use of a properly arranged video camera native to the device running the app, adding another possible barrier to their use. To our knowledge, no free VJ apps relying on the built-in accelerometer of a smart phone have been tested. The ability to use a free app that requires no outside help or special set-up could broadly increase use of the VJ test in various clinical and non-clinical settings.

The purpose of this study was to assess the validity and reliability of the free Jumpster app (Skyhawk Media, LLC, Palmdale, CA) by comparing the VJ height results obtained during a countermovement jump (CMJ) to that of the Vertec. Based on pilot data, we hypothesized that the Jumpster app would provide moderate validity and reliability when compared to the Vertec.

## Materials and methods

Our target sample size was determined using data from a similarly designed study published by Yingling et al. (Yingling et al., 2018). Analysis of this data in JMP (version 17.1; SAS Institute Inc., Cary, NC) yielded a calculated power of approximately 80% with a sample size of 20 participants, while a sample size of 40 participants resulted in nearly 99% power. Given these results the study team aimed to recruit 40 participants from the University student population via social media and word of mouth between February and March of 2024. Apparently healthy men and women between the ages of 18 – 50 years were recruited for participation. Among those, individuals reporting any known medical condition that would preclude jumping activities, such as those who sustained a lower extremity injury in the past 6 months or those with a history of neurologic or lower extremity arthritic conditions, were excluded from engaging in the study. Each participant was informed of the purposes and requirements of the study prior to providing the study team with their written informed consent. All study methodologies were reviewed and approved by the University Institutional Review Board (IRB00004838FWA00008717IORG0004078).

Following consent and prior to testing, the study team assessed each participant’s height and weight, and captured basic demographic information. Next, participants were instructed on how to complete a standardized CMJ technique using the Vertec as previously outlined by Yingling et al. (Yingling et al., 2018). Following instruction, each participant’s baseline vertical reach was determined, and they were fitted with a special belt harness to hold an iPhone SE3 loaded with the Jumpster app. While the app developer suggested placing the phone in each participant’s pocket, early pilot testing indicated this was not sufficient to secure the device for all participants. Thus, the addition of a harness provided a more secure method of holding the device to reduce excess movement during testing. Participants were instructed to execute at least 3 CMJ trials with 1-minute of rest between each jump. In instances where the Jumpster app did not record data, up to 2 additionally trials were permitted; however, if all original 3 trials were unsuccessful then participants were dismissed, and their data would be removed for analysis. Just prior to the beginning of each trial, both the app and the Vertec were reset by the study team and basic jumping instructions provided again. Following each trial, data from both devices was recorded by a single member of the study team. No feedback on jump height or overall jump performance was provided to participants during testing.

The Vertec data was measured in inches and converted to centimetres, with VJ height calculated by subtracting the baseline reach height from the measured jump height for each trial. The Jumpster app automatically provides users with a calculated VJ height in inches which was also converted into centimetres; no additional calculations were necessary by the study team for the Jumpster app data. An overall failure rate for the mobile app (i.e., trials where no usable data was obtained) was also calculated across all participants and trials and presented as a percentage (total failed trials over total trials).

Only device-matched trials were used for analysis, while trials with missing data for either measurement tool were discarded entirely. Data for both measurement tools were independently assessed for normality using the Shapiro-Wilk test. Next, descriptive statistics were calculated for participant age, height, weight, maximum Vertec jump height, and maximum Jumpster jump height. Independent samples Welch’s t-tests were also conducted for these variables to assess possible differences between men and women.

As previously outlined, several analysis tools are helpful in determining the validity and reliability of different measures or devices (Giavarina, 2015; Van Stralen et al., 2012). Our assessment of validity began by measuring the relationship between the Vertec and Jumpster tools using a correlation analysis and visually comparing the line of equality to the calculated regression line. The relative orientation of both lines provides additional insights into the validity of any significant relationship. Next, a tolerance analysis (similar to a Bland-Altman analysis) accounting for intraparticipant correlations in the model structure was conducted to assess possible measurement bias between the two tools using the SimplyAgree package (version 0.2.0) in RStudio (version 2024.04.1+748; Posit Software, PBC, Boston, MA) (Caldwell, 2022; Francq et al., 2020). To provide contextually-relevant absolute bounds of measurement error we also established acceptable limits of variability based on our calculated standard error of the measure (SEM) for the Vertec data (outlined below) and overlaid these results on a Bland-Altman styled plot as another means of assessing overall tool validity (Giavarina, 2015).

Reliability was first assessed in absolute terms by calculating the 95% predictive intervals (PI_95_) and tolerance limits (TL_95_) between the two measures (similar to Bland-Altman limits of agreement±95% CI) (Caldwell, 2022; Francq et al., 2020). Additional assessments of absolute reliability were also conducted by calculating the SEM and coefficient of variation (CV) for both tools using mean square error estimations, again with the SimplyAgree package (CITATION: Caldwell, 2022) (Weir, 2005). Finally, relative reliability was assessed by estimating the intraclass correlation coefficient and 95% confidence intervals between the Vertec and Jumpster app, based on a 2-way random effects, mean rating, absolute agreement model (ICC 2,K) (CITATION: Caldwell, 2022) (Koo & Li, 2016; Weir, 2005). This was also calculated using mean square error estimations in the SimplyAgree package. All data were analyzed using RStudio (α = 0.05).

## Results

In total 38 participants were recruited, all of whom were able to complete testing without difficulty or insult. However, the Jumpster app failed across all trials for 2 participants resulting in their data being removed entirely for analysis. Among the remaining 36 participants, an additional 13 app failures were logged resulting trial data across 9 individuals also being removed for analysis. An overall app failure rate of 15.97% was catalogued. Despite multiple app failures, members of the study team only provided 3 participants with an additional fourth trial and 1 participant with 2 additional trials (a fourth and fifth trial). Even with this inconsistency, all participants successfully completed on average 2.8 trials with both tools simultaneously (at least 2 trials per individual), resulting in 100 successful matched trials for analysis. These data were normally distributed, and analysis proceeded as planned. As outlined in Table 1, when compared to women, men were significantly taller (p < 0.01) and heavier (p < 0.01); however, both groups were similarly aged (p = 0.06). Additionally, men were able to achieve significantly greater maximum jump heights when measured using the Vertec (see Table 1; p <0.01); however, no such differences were observed with the Jumpster app (p = 0.25). The lack of a statistically significant difference in maximal jump height between men and women for the Jumpster app only, could indicate poor validity of the device.

**Table 1.** Summary data for all subjects.

|  | All | Men | Women | Men vs Women |
| --- | --- | --- | --- | --- |
| N | 36 | 22 | 14 |  |
| Age (years) | 22.7 ± 6.4 | 24.0 ± 7.9 | 20.6 ± 1.0 | p = 0.06 |
| Height (cm) | 172.7 ± 9.2 | 178.1 ± 6.0 | 164.3 ± 6.7 | p < 0.01 |
| Weight (kg) | 81.4 ± 18.8 | 90.9 ± 16.3 | 66.4 ± 11.4 | p < 0.01 |
| Max Vertec Jump Height (cm) | 51.5 ± 11.3 | 56.7 ± 9.1 | 43.4 ± 9.8 | p < 0.01 |
| Max Jumpster Jump Height (cm) | 47.0 ± 16.5 | 49.3 ± 19.4 | 43.5 ± 10.0 | p = 0.25 |

A significant, weak and positive relationship was observed between the two measurement tools (r = 0.24, p = 0.01; see Figure 1). Additionally, the regression line fell crossed the line of equality likely indicating a proportional bias (Coskun, 2024). A significant, moderately strong and positive relationship was in fact observed when the data were fitted to a Bland-Altman styled plot (see Figure 2; r = 0.45; p < 0.01); resulting in the tolerance model being adjusted for a proportional bias. The resulting proportional bias had a slope [±95% CI] of 1.59 [±0.06] and intercept [±95% CI] of -80.73 [±6.94] cm. When paired with the calculated acceptable limits of variability for the Vertec (±3.57 cm), the Jumpster app generally underestimated jump height when compared to the Vertec below a height of ∼ 48.0 – 53.0 cm and overestimated jump height beyond that range. Weak correlation between the measures paired with comparisons of our assessed proportional bias to the Vertec limits of variability generally indicates poor validity of the Jumpster app.

**Figure 1.**
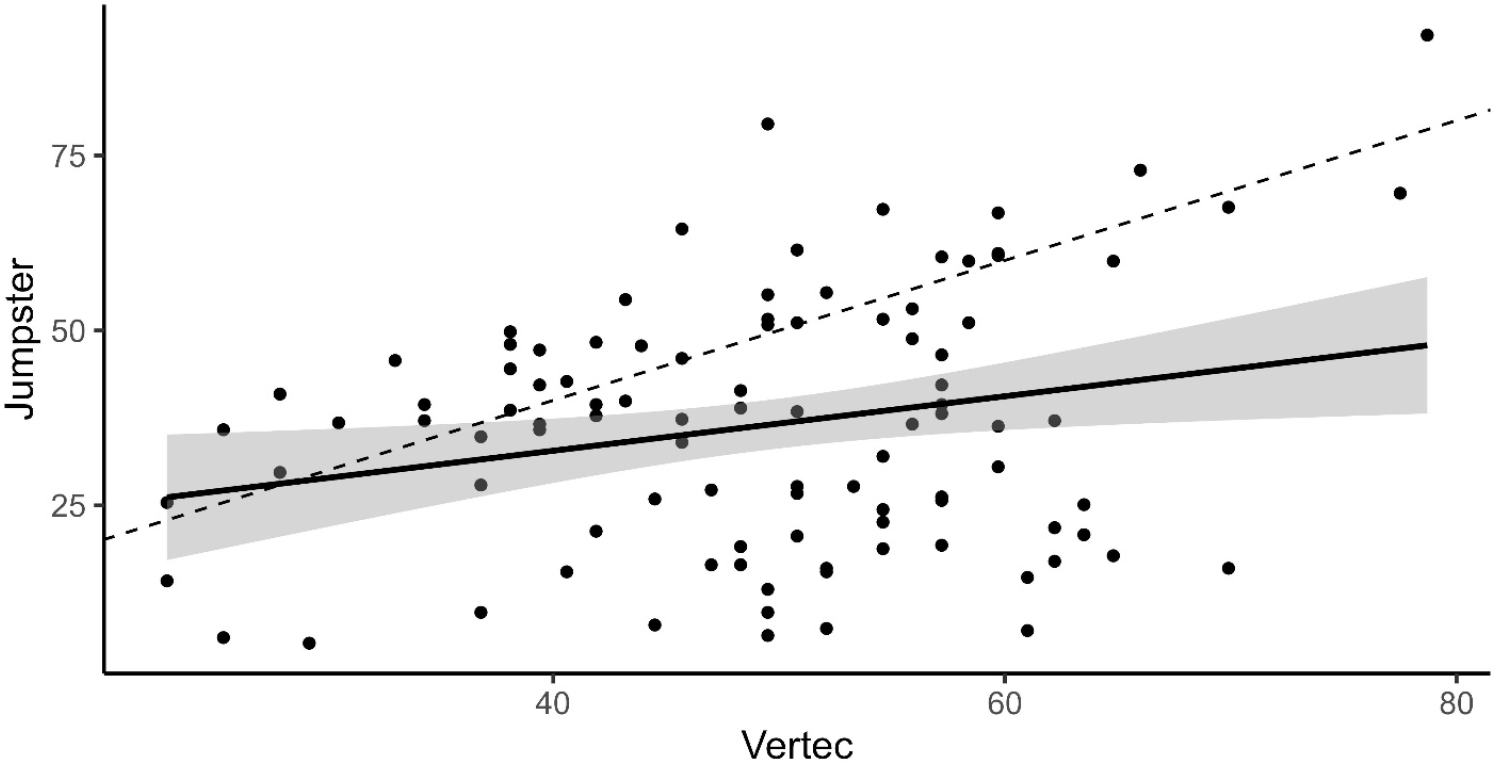
Regression line (solid), calculated 95% CI of the regression line (grey), and the line of equality (dashed) overlaid a scatterplot of all matched trials.

**Figure 2.**
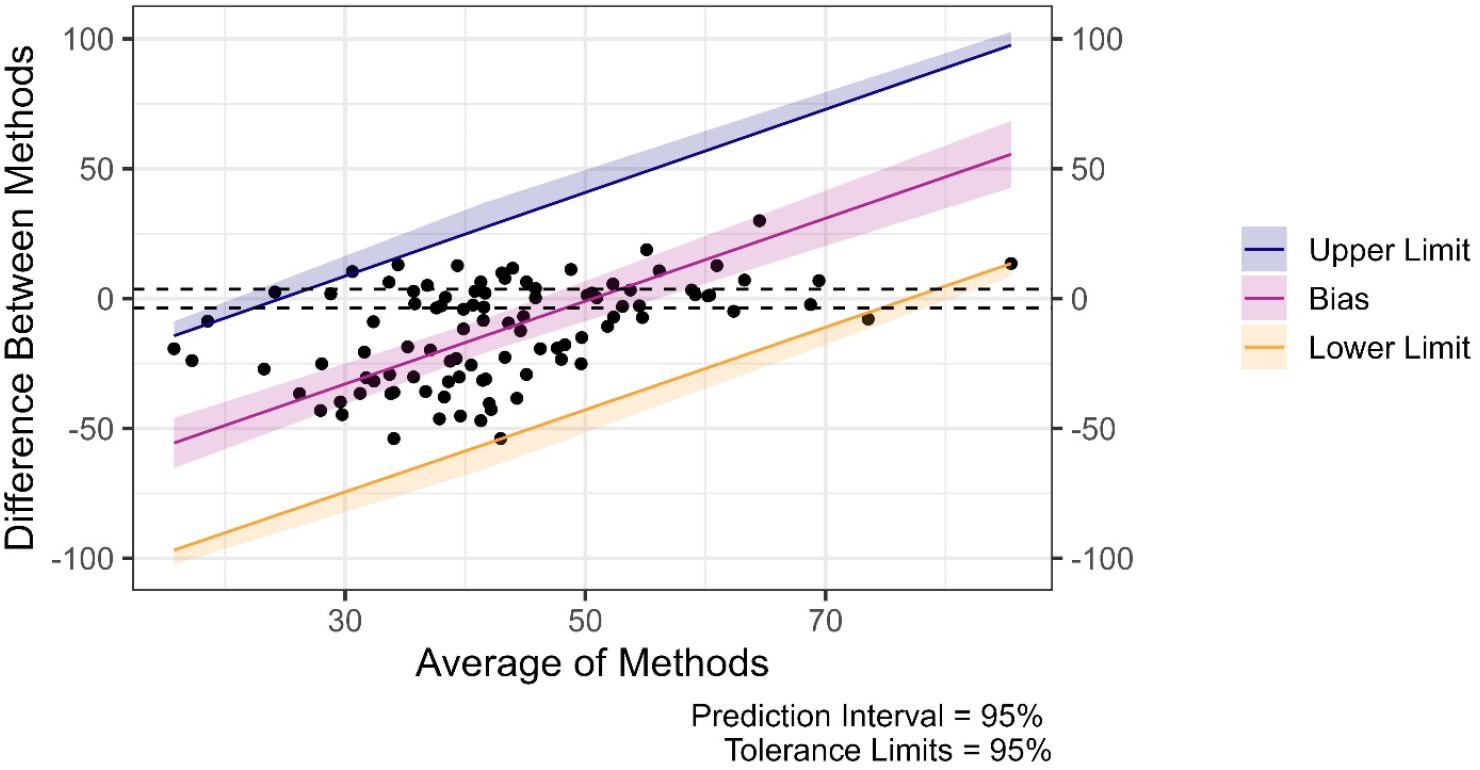
Bland-Altman styled plot with proportional bias ± 95% CI (red), both upper (blue) and lower (yellow) 95% predictive intervals and tolerance limits, and estimated limits of variability for the Vertec (dashed lines) depicted. X-axis: average of jump heights using both the Jumpster and the Vertec; y-axis: difference in jump heights between both tools (Jumpster – Vertec).

Potentially indicating good reliability of the Jumpster tool, all plotted datapoints on our Bland-Altman styled plot fell within the calculated PI_95_[±TL_95_] (upper: slope:1.61[+0.02], intercept: -39.54[+7.49] cm; lower: slope:1.58[-0.02], intercept: -21.92[-7.49] cm). However, the SEM and CV of the Vertec (SEM: 3.57 cm; CV: 7.22%) were both smaller than the Jumpster (SEM: 14.7 cm; CV: 40.30%), indicating likely poor reliability of the Jumpster app. Also indicating poor reliability of the Jumpster app, the estimated ICC [upper and lower 95% CI] between the two devices was assessed to be 0.55 [0.79, - 0.08].

## Discussion

To our knowledge this is the first study to examine whether the Jumpster VJ app, which relies on the built-in accelerometer of a smart phone, provides comparable results to the Vertec VJ test system. Based on prior pilot data, we hypothesized that the Jumpster app would provide moderate reliability and validity. Unfortunately with an overall failure rate just over 15% and significantly greater internal and external inconsistency (when compared to the Vertec), our results indicate that the app is neither reliable or valid.

While there is a continuing need in sports science to develop innovative, simple, and affordable tools to measure performance, it is essential that such devices produce are highly reliable and at least acceptably valid. A significant body of support exists for movement scientists to endorse the use of various video based VJ apps. (Bogataj, Pajek, Andrašić, et al., 2020; Cruvinel-Cabral et al., 2018; Gallardo-Fuentes et al., 2016; Gençoğlu et al., 2023; Oliveira-Silva et al., 2018; Stanton et al., 2015; Yingling et al., 2018). The relative “gold standard” among this class of VJ apps appears to be *My Jump 2*, which was developed more than a decade ago. This app relies on positioning a smart phone, ipad, or Mac compute to record live video with a native camera, allowing users to ascertain flight time which is then used to calculate jump height (Gençoğlu et al., 2023). In a recent meta-analysis, Gencoglu et al. (Gençoğlu et al., 2023) synthesized 21 studies consisting of 839 participants and revealed a high level of agreement between the My Jump app and criterion measures with near perfect reliability. More recent data using the What’s my Vertical and Jumpo apps have also shown good validity and high reliability. In contrast, the Jumpster app relies on indwelling accelerometers and is advertised as an accurate and easy to use tool, allowing one to simply slip their phone in their pocket and measure their VJ. Our study was designed to test the app under standardized conditions and much more stringent controls than are typically encountered in the field. Despite these efforts, the app failed to produce either reliable or valid results.

It was noted during pilot testing that placing the phone in a pocket could produce excess movement (even sometimes resulting in the phone involuntarily exiting the user’s pocket), so a more uniform restraint was employed, which itself needed to be further secured after some initial test failures. Even after minimizing these extra movements of the device, the app still proved unsuccessful in producing any result several times. The initial question of whether the accelerometers in the iPhone are valid and reliable was posed. Smart phone based accelerometers have been shown to be reliable and valid for walking (Strongman et al., 2023), running (Balsalobre-Fernández et al., 2017), weight lifting (Peláez Barrajón & San Juan, 2020), and even jumping (Mateos-Angulo et al., 2020) activities. In the latter study, Mateos-Angulo et al. (Mateos-Angulo et al., 2020) used the xSensor Pro app (no longer available) to obtain direct acceleration data to then calculate jump time and height, comparable to the My Jump 2 app. Similarly, a study examining the bench press noted only moderate reliability and validity when measuring lifting velocity, which was still far better than we were able to achieve when assessing VJ height during a CMJ. The authors, however, point to a possible explanation for our poor results, noting that assessment of the data by an app is highly dependent on the calibration of the device and algorithm used to interpret it. Thus, we surmise that the iPhone hardware likely captures the target data accurately, but the Jumpster app fails to accurately or reliably filter “noise” and/or interpret the data correctly.

## Conclusions

Compared to the Vertec VJ measurement tool, the iPhone accelerometer based Jumpster app failed to produce reliable or valid results in a cohort of college aged men and women. The authors recommend that individuals and researchers rely on the more proven video-based analysis tools for VJ assessment if a moblie app is desired over more traditional tools like the Vertec. More research and development are needed on the application of accelerometer-based apps for sports performance.

## Acknowledgements

We would like to acknowledge both John Ward and Brenna Kehoe for their help in collecting our data and compiling the literature for this research. We would also like to thank Dr. Aaron Caldwell for his correspondence and assistance with using the SimplyAgree package in R.

## Declaration of Interests

Both authors served as participants in this study; however, they were not involved with their own data collection or made aware of their results until data processing. No other potential conflicts of interest are reported by the authors.

## Data Availability

The data that support the findings of this study are available from the corresponding author, upon reasonable request.

## Notes

### Competing Interest Statement

The authors have declared no competing interest.

## References

Alcazar, J., Aagaard, P., Haddock, B., Kamper, R. S., Hansen, S. K., Prescott, E., Alegre, L. M., Frandsen, U., & Suetta, C. (2020). Age- and Sex-Specific Changes in Lower-Limb Muscle Power Throughout the Lifespan. The Journals of Gerontology: Series A, 75(7), 1369–1378. 10.1093/gerona/glaa013

Argaud, S., Pairot De Fontenay, B., Blache, Y., & Monteil, K. (2017). Explosive movement in the older men: Analysis and comparative study of vertical jump. Aging Clinical and Experimental Research, 29(5), 985–992. 10.1007/s40520-016-0660-0

Balsalobre-Fernández, C., Agopyan, H., & Morin, J.-B. (2017). The Validity and Reliability of an iPhone App for Measuring Running Mechanics. Journal of Applied Biomechanics, 33(3), 222–226. 10.1123/jab.2016-0104

Baptista, F., Zymbal, V., & Janz, K. F. (2022). Predictive Validity of Handgrip Strength, Vertical Jump Power, and Plank Time in the Identification of Pediatric Sarcopenia. Measurement in Physical Education and Exercise Science, 26(4), 361–370. 10.1080/1091367X.2021.1987242

Bogataj, Š., Pajek, M., Andrašić, S., & Trajković, N. (2020). Concurrent validity and reliability of my jump 2 app for measuring vertical jump height in recreationally active adults. Applied Sciences, 10(11), 3805.

Bogataj, Š., Pajek, M., Hadžić, V., Andrašić, S., Padulo, J., & Trajković, N. (2020). Validity, Reliability, and Usefulness of My Jump 2 App for Measuring Vertical Jump in Primary School Children. International Journal of Environmental Research and Public Health, 17(10), 3708. 10.3390/ijerph17103708

Buckthorpe, M., Morris, J., & Folland, J. P. (2012). Validity of vertical jump measurement devices. Journal of Sports Sciences, 30(1), Article 1. 10.1080/02640414.2011.624539

Caldwell, A. R. (2022). SimplyAgree: An R package and jamovi Module forSimplifying Agreement and Reliability Analyses. Journal of Open Source Software, 7(71), 4148. 10.21105/joss.04148

Coskun, A. (2024). Bias in Laboratory Medicine: The Dark Side of the Moon. Annals of Laboratory Medicine, 44(1), 6–20. 10.3343/alm.2024.44.1.6

Cruvinel-Cabral, R. M., Oliveira-Silva, I., Medeiros, A. R., Claudino, J. G., Jiménez-Reyes, P., & Boullosa, D. A. (2018). The validity and reliability of the “My Jump App” for measuring jump height of the elderly. PeerJ, 6, e5804.

Dias, J. A., Pupo, J. D., Reis, D. C., Borges, L., Santos, S. G., Moro, A. R., & Borges, N. G. (2011). Validity of Two Methods for Estimation of Vertical Jump Height. Journal of Strength and Conditioning Research, 25(7), 2034–2039. 10.1519/JSC.0b013e3181e73f6e

Dow, M., Aguinaldo, A., Matthews, J., & Ogren, A. (n.d.). Concurrent validity and reliability of mobile applications in measuring vertical jump performance.

Faigenbaum, A. D., & MacDonald, J. P. (2017). Dynapenia: It’s not just for grown-ups anymore. Acta Paediatrica, 106(5), 696–697. 10.1111/apa.13797

Faigenbaum, A. D., Ratamess, N. A., Kang, J., Bush, J. A., & Rebullido, T. R. (2023). May the Force Be with Youth: Foundational Strength for Lifelong Development. Current Sports Medicine Reports, 22(12), 414–422.

Forte, R., & Macaluso, A. (2008). Relationship between performance-based and laboratory tests for lower-limb muscle strength and power assessment in healthy older women. Journal of Sports Sciences, 26(13), 1431–1436. 10.1080/02640410802208994

Francq, B. G., Berger, M., & Boachie, C. (2020). To tolerate or to agree: A tutorial on tolerance intervals in method comparison studies with BivRegBLS R Package. Statistics in Medicine, 39(28), 4334–4349. 10.1002/sim.8709

Gallardo-Fuentes, F., Gallardo-Fuentes, J., Ramírez-Campillo, R., Balsalobre-Fernández, C., Martínez, C., Caniuqueo, A., Cañas, R., Banzer, W., Loturco, I., Nakamura, F. Y., & Izquierdo, M. (2016). Intersession and Intrasession Reliability and Validity of the My Jump App for Measuring Different Jump Actions in Trained Male and Female Athletes. Journal of Strength and Conditioning Research, 30(7), Article 7. 10.1519/JSC.0000000000001304

Gençoğlu, C., Ulupınar, S., Özbay, S., Turan, M., SavaŞ, B. Ç., Asan, S., & Ince, I. (2023). Validity and reliability of “My Jump app” to assess vertical jump performance: A meta-analytic review. Scientific Reports, 13(1), Article 1. 10.1038/s41598-023-46935-x

Giavarina, D. (2015). Understanding Bland Altman analysis. Biochemia Medica, 25(2), 141–151. 10.11613/BM.2015.015

Katartzi, E., Gantiraga, E., & Komsis, G. (2005). The relationship between specific strength components of lovver limbs and vertical jumping ability in school-aged children. J. Human Movement Stud, 48, 227–243.

Koo, T. K., & Li, M. Y. (2016). A Guideline of Selecting and Reporting Intraclass Correlation Coefficients for Reliability Research. Journal of Chiropractic Medicine, 15(2), 155–163. 10.1016/j.jcm.2016.02.012

Leard, J. S., Cirillo, M. A., Katsnelson, E., Kimiatek, D. A., Miller, T. W., Trebincevic, K., & Garbalosa, J. C. (2007). Validity of two alternative systems for measuring vertical jump height. The Journal of Strength & Conditioning Research, 21(4), 1296–1299.

Mateos-Angulo, A., Galán-Mercant, A., & Cuesta-Vargas, A. I. (2020). Kinematic Mobile Drop Jump Analysis at Different Heights Based on a Smartphone Inertial Sensor. Journal of Human Kinetics, 73(1), 57–65. 10.2478/hukin-2019-0131

Montalvo, S., Gonzalez, M. P., Dietze-Hermosa, M. S., Eggleston, J. D., & Dorgo, S. (2021). Common Vertical Jump and Reactive Strength Index Measuring Devices: A Validity and Reliability Analysis. Journal of Strength and Conditioning Research, 35(5), 1234–1243. 10.1519/JSC.0000000000003988

Oh, D.-S., Choi, Y.-H., Shim, Y.-J., Park, S.-H., & Lee, M.-M. (2020). Concurrent validity, inter-, and intrarater reliabilities of smart device based application for measuring vertical jump performance. Baltic Journal of Health and Physical Activity, 12(3), 4.

Oliveira-Silva, I., Cruvinel-Cabral, R. M., Medeiros, A. R., & Boullosa, D. A. (2018). Validity of My Jump App to Measure Vertical Jump Height of the Elderly: 2484 Board #8 June 1 1 00 PM - 3 00 PM. Medicine & Science in Sports & Exercise, 50(5S), Article 5S. 10.1249/01.mss.0000537112.08878.12

Peláez Barrajón, J., & San Juan, A. F. (2020). Validity and Reliability of a Smartphone Accelerometer for Measuring Lift Velocity in Bench-Press Exercises. Sustainability, 12(6), 2312. 10.3390/su12062312

Pupo, J. D., Ache-Dias, J., Kons, R. L., & Detanico, D. (2020). Are vertical jump height and power output correlated to physical performance in different sports? An allometric approach. Human Movement, 22(2), Article 2. 10.5114/hm.2021.100014

Runge, M., Rittweger, J., Russo, C. R., Schiessl, H., & Felsenberg, D. (2004). Is muscle power output a key factor in the age-related decline in physical performance? A comparison of muscle cross section, chair-rising test and jumping power. Clinical Physiology and Functional Imaging, 24(6), 335–340. 10.1111/j.1475-097X.2004.00567.x

Santos, C. A. F., Amirato, G. R., Jacinto, A. F., Pedrosa, A. V., Caldo-Silva, A., Sampaio, A. R., Pimenta, N., Santos, J. M. B., Pochini, A., & Bachi, A. L. L. (2022). Vertical Jump Tests: A Safe Instrument to Improve the Accuracy of the Functional Capacity Assessment in Robust Older Women. Healthcare, 10(2), 323. 10.3390/healthcare10020323

Shur, N. F., Creedon, L., Skirrow, S., Atherton, P. J., MacDonald, I. A., Lund, J., & Greenhaff, P. L. (2021). Age-related changes in muscle architecture and metabolism in humans: The likely contribution of physical inactivity to age-related functional decline. Ageing Research Reviews, 68, 101344. 10.1016/j.arr.2021.101344

Stanton, R., Kean, C. O., & Scanlan, A. T. (2015). My Jump for vertical jump assessment. British Journal of Sports Medicine, 49(17), Article 17. 10.1136/bjsports-2015-094831

Stojanović, D., Savić, Z., Vidaković, H. M., Stojanović, T., Momčilović, Z., & Stojanović, T. (2020). RELATIONSHIP BETWEEN BODY COMPOSITION AND VERTICAL JUMP PERFORMANCE AMONG ADOLESCENTS. Acta Medica Medianae, 59(1), Article 1. 10.5633/amm.2020.0109

Strongman, C., Cavallerio, F., Timmis, M. A., & Morrison, A. (2023). A Scoping Review of the Validity and Reliability of Smartphone Accelerometers When Collecting Kinematic Gait Data. Sensors, 23(20), 8615. 10.3390/s23208615

Temfemo, A., Hugues, J., Chardon, K., Mandengue, S.-H., & Ahmaidi, S. (2009). Relationship between vertical jumping performance and anthropometric characteristics during growth in boys and girls. European Journal of Pediatrics, 168(4), 457–464. 10.1007/s00431-008-0771-5

Trombetti, A., Reid, K. F., Hars, M., Herrmann, F. R., Pasha, E., Phillips, E. M., & Fielding, R. A. (2016). Age-associated declines in muscle mass, strength, power, and physical performance: Impact on fear of falling and quality of life. Osteoporosis International, 27(2), 463–471. 10.1007/s00198-015-3236-5

Van Stralen, K. J., Dekker, F. W., Zoccali, C., & Jager, K. J. (2012). Measuring Agreement, More Complicated Than It Seems. Nephron Clinical Practice, 120(3), c162–c167. 10.1159/000337798

Vieira, A., Blazevich, A. J., DA Costa, A. S., Tufano, J. J., & Bottaro, M. (2021). Validity and Test-retest Reliability of the Jumpo App for Jump Performance Measurement. International Journal of Exercise Science, 14(7), 677–686.

Vieira, A., Ribeiro, G. L., Macedo, V., De Araújo Rocha Junior, V., Baptista, R. D. S., Gonçalves, C., Cunha, R., & Tufano, J. (2023). Evidence of validity and reliability of Jumpo 2 and MyJump 2 for estimating vertical jump variables. PeerJ, 11, e14558. 10.7717/peerj.14558

Weir, J. P. (2005). Quantifying Test-Retest Reliability Using the Intraclass Correlation Coefficient and the SEM. The Journal of Strength and Conditioning Research, 19(1), 231. 10.1519/15184.1

Yingling, V. R., Castro, D. A., Duong, J. T., Malpartida, F. J., Usher, J. R., & O, J. (2018). The reliability of vertical jump tests between the Vertec and My Jump phone application. PeerJ, 6, e4669. 10.7717/peerj.4669

